# Rapid metagenomic identification of viral pathogens in clinical samples by real-time nanopore sequencing analysis

**DOI:** 10.1101/020420

**Authors:** Alexander L. Greninger, Samia N. Naccache, Scot Federman, Guixia Yu, Placide Mbala, Vanessa Bres, Doug Stryke, Jerome Bouquet, Sneha Somasekar, Jeffrey M. Linnen, Roger Dodd, Prime Mulembakani, Bradley S. Schneider, Jean-Jacques Muyembe, Susan L. Stramer, Charles Y. Chiu

**Author notes:** These authors contributed equally to this work (Samia N. Naccache, Scot Federman).

## Abstract

We report unbiased metagenomic detection of chikungunya virus (CHIKV), Ebola virus (EBOV), and hepatitis C virus (HCV) from four human blood samples by MinION nanopore sequencing coupled to a newly developed, web-based pipeline for real-time bioinformatics analysis on a computational server or laptop (MetaPORE). At titers ranging from 10^7^-10^8^ copies per milliliter, reads to EBOV from two patients with acute hemorrhagic fever and CHIKV from an asymptomatic blood donor were detected within 4 to 10 minutes of data acquisition, while lower titer HCV virus (1x10^5^ copies per milliliter) was detected within 40 minutes. Analysis of mapped nanopore reads alone, despite an average individual error rate of 24% [range 8-49%], permitted identification of the correct viral strain in all 4 isolates, and 90% of the genome of CHIKV was recovered with >98% accuracy. Using nanopore sequencing, metagenomic detection of viral pathogens directly from clinical samples was performed within an unprecedented <6 hours sample-to-answer turnaround time and in a timeframe amenable for actionable clinical and public health diagnostics.

## Background

Acute febrile illness has a broad differential diagnosis and can be caused by a variety of pathogens. Metagenomic next-generation sequencing (NGS) is particularly attractive for diagnosis and public health surveillance of febrile illness because the approach can broadly detect viruses, bacteria, and parasites in clinical samples by uniquely identifying sequence data [1, 2]. Although currently limited by a sample-to-answer turnaround time of >20 hr (Fig. 1), we and others have reported that unbiased pathogen detection using metagenomic NGS can generate actionable results in timeframes relevant to clinical diagnostics [3-6] and public health [7, 8]. As sequence reads are generated in parallel and not in series, real-time analysis by second-generation platforms such as Illumina and Ion Torrent has been hampered by the need to wait until a sufficient read length has been achieved for diagnostic pathogen identification.

**Figure 1.**
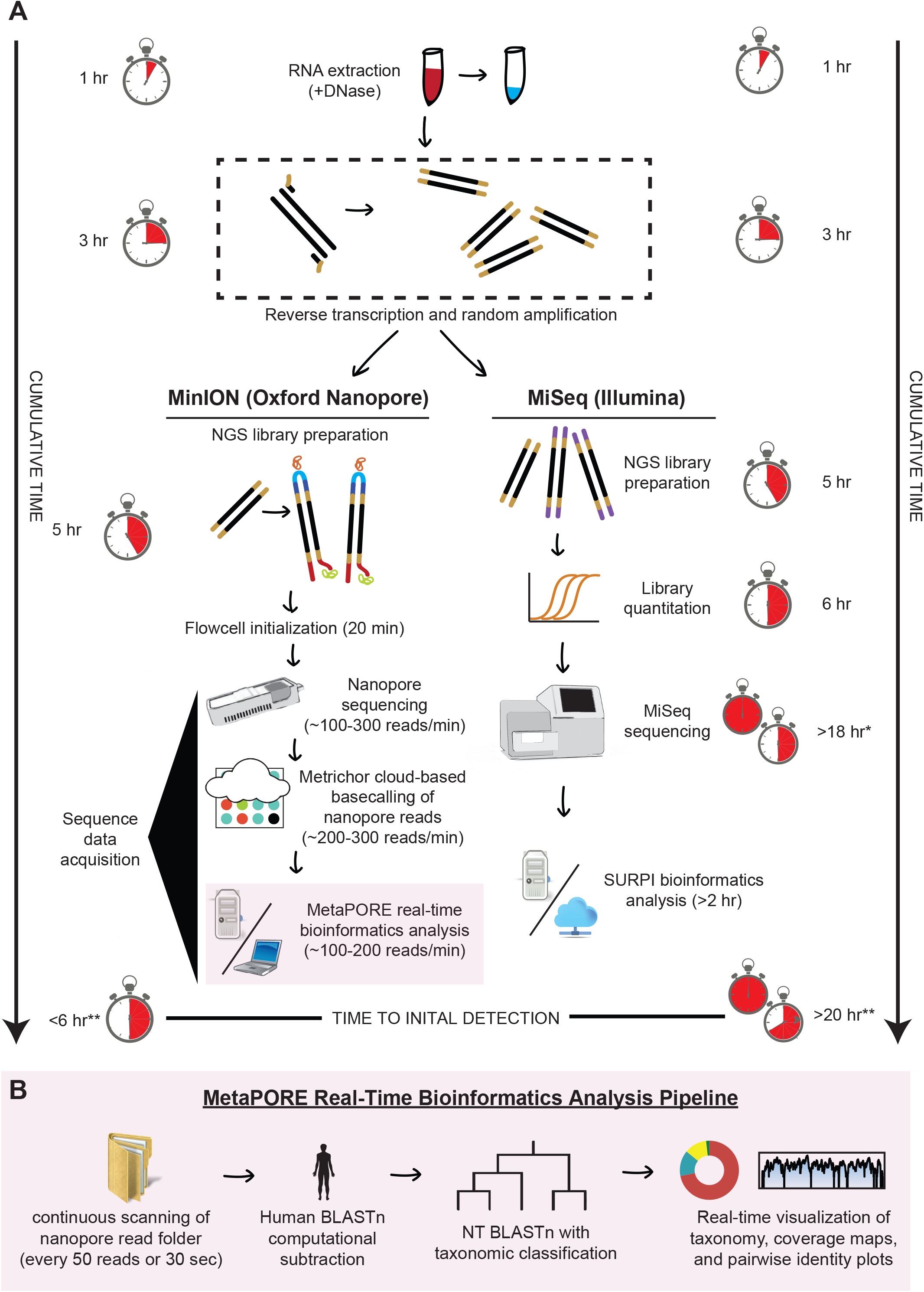
Metagenomic sequencing workflow for MinION nanopore sequencing compared to Illumina MiSeq sequencing. The turnaround time for sample-to-detection nanopore sequencing, defined here as the cumulative time taken for nucleic acid extraction, reverse transcription, library preparation, sequencing, MetaPORE bioinformatics analysis, and detection, was under 6 hr, while Illumina sequencing took over 20 hr. The time differential is accounted for by increased times for library quantitation, sequencing, and bioinformatics analysis with the Illumina protocol. *assumes a 12-hr 50 nucleotide (nt) single-end MiSeq run of ∼12-15 million reads, with 50 nt the minimum read length needed for accurate pathogen identification. **denotes estimated average SURPI bioinformatics analysis run length for MiSeq data[15]. The stopwatch is depicted as a 12-hr clock.

Nanopore sequencing is a third-generation sequencing technology that has two key advantages over second-generation technologies – longer reads and the ability to perform real-time sequence analysis. To date, the longer nanopore reads have enabled scaffolding of prokaryotic and eukaryotic genomes and sequencing of bacterial and viral cultured isolates [9-12], but the platform’s capacity for real-time metagenomic analysis of clinical samples has not yet been leveraged. As of mid-2015, the MinION nanopore sequencer is capable of producing at least 100,000 sequences with an average read length of 5 kB, in total producing up to 1 Gb of sequence in 24 hours on one flow cell [13]. Here we present using nanopore sequencing for metagenomic detection of viral pathogens from clinical samples with sample-to-answer turnaround times of under 6 hours. We also present MetaPORE, a real-time, web-based sequence analysis and visualization tool for pathogen identification from nanopore data.

## Materials and Methods

### MAP program

Since July 2014, our lab has participated in the MinION Access Program (MAP), an early access program for beta users of the Oxford Nanopore MinION. Program participants receive free flow cells and library preparation kits for testing and validation of new protocols and applications on the MinION platform. During our time in the MAP program, we have seen significant progress in quality control and sequencing yield of flow cells (Table 1).

### Nucleic acid extraction

Total nucleic acid was extracted from 400 μL of a chikungunya virus (CHIKV)-positive serum sample previously collected from an asymptomatic blood donor during the 2014 CHIKV outbreak in Puerto Rico (Chik1) [14]. The serum sample was inactivated in a 1:3 ratio of TRIzol LS (Life Technologies, Carlsbad, CA, USA) at the American Red Cross prior to shipping to University of California, San Francisco (UCSF). Direct-zol RNA MiniPrep (Zymo Research, Irvine, CA, USA) was used for nucleic acid extraction, including on-column treatment with Turbo DNAse (Life Technologies) for 30 min at 37°C to deplete human host genomic DNA.

For the Ebola virus (EBOV) specimens, total nucleic acid was extracted using a QIAamp Viral RNA kit (Qiagen, Valencia, CA, USA) from 140 μL of whole blood from two patients with suspected Ebola hemorrhagic fever from a 2014 outbreak in the Democratic Republic of the Congo (DRC) (Ebola1 and Ebola2). RNA was extracted at Institut National de Recherche Biomédicale in Kinshasa, DRC, preserved using RNAstable (Biomatrica, San Diego, CA, USA), and shipped at room temperature to UCSF. Upon receipt, the extracted RNA sample was treated with 1 μL Turbo DNase (Life Technologies), followed by clean-up using the Direct-zol RNA MiniPrep Kit (Zymo Research).

For the HCV sample, an HCV-positive serum specimen at a titer of 1.6x10^7^ cp/mL (HepC1) was diluted to 1x10^5^ cp/mL using pooled negative serum. Total nucleic acid was then extracted from 400uL of serum using the EZ1 Viral RNA kit followed by treatment with Turbo DNase at 30 min at 37°C and clean-up using the RNA Clean and Concentrator Kit (Zymo Research).

### Molecular confirmation of viral infection

A previously reported TaqMan quantitative reverse-transcription PCR (qRT-PCR) assay targeting the EBOV NP gene was used for detection of EBOV and determination of viral load. The assay was run on a Stratagene MX300P real-time PCR instrument and performed using the TaqMan Fast Virus 1-Step Master Mix (Life Technologies) in 20 μL total reaction volume (5 μL 4x Taqman mix, 1ul sample extract), with 0.75uM of each primer (F565 5’-TCTGACATGGATTACCACAAGATC-3’, R640 5’- GGATGACTCTTTGCCGAACAATC-3’) and 0.6uM of the probe (p597S 6FAM-AGGTCTGTCCGTTCAA-MGBNFQ). Conditions for the qRT-PCR were as follows: 50°C, 10minutes / 95°C, 20 s followed by 45 cycles of 95°C 3s / 60°C, 30 s. Viral copy number was calculated by standard curve analysis with a plasmid vector containing the EBOV amplicon. The first EBOV sample analyzed by nanopore sequencing (Ebola1) corresponded to the Ebola virus/*H.sapiens*- wt/COD/2014/Lomela-Lokolia16 strain, while the second Ebola sample (Ebola2) corresponded to the Ebola virus/*H.sapiens*-wt/COD/2014/Lomela-LokoliaB11 strain. The CHIKV-positive sample was identified and quantified using a transcription-mediated amplification (TMA) assay (Hologic, Bedford, MA, USA) as previously described [14]. HCV was quantified using the FDA-approved Abbott RealTi*m*e RT-PCR assay as performed in the UCSF Clinical Microbiology Laboratory on the Abbott Molecular m2000 system.

### Construction of metagenomic amplified cDNA libraries

To obtain ≥1 μg of metagenomic cDNA library required for the nanopore sequencing protocol, randomly amplified cDNA was generated using a primer-extension pre-amplification method (Round A/B) as described previously [15-17]. Of note, this protocol has been extensively tested on clinical samples for metagenomic pan-pathogen detection of DNA and RNA viruses, bacteria, fungi, and parasites [4, 6, 15, 17, 18]. Briefly, in Round A, RNA was reverse-transcribed with SuperScript III Reverse Transcriptase (Life Technologies,) using Sol-PrimerA (5’-GTTTCCCACTGGAGGATA-N_9_-3’), followed by second-strand DNA synthesis with Sequenase DNA polymerase (Affymetrix, Santa Clara, CA, USA). Reaction conditions for Round A were as follows: 1 μL of Sol-PrimerA (40 pmol/μl) was added to 4 μl of sample RNA, heated at 65°C for 5 minutes, then cooled at room temperature for 5 minutes. Five μL of SuperScript Master Mix (2 μl 5X First-Strand Buffer, 1 μL water, 1 μL 12.5 mM dNTP mix, 0.5 μL 0.1M DTT, 0.5 μL SS III RT) was then added and incubated at 42°C for 60 minutes. For second strand synthesis, 5 μL of Sequenase Mix #1 (1 μL 5X Sequenase Buffer, 3.85 μL ddH_2_O, 0.15 μL Sequenase enzyme) was added to the reaction mix and incubated at 37°C x 8 min, followed by addition of Sequenase Mix #2 (0.45 μl Sequenase Dilution Buffer, 0.15 μl Sequenase Enzyme) and a second incubation at 37°C x 8 min. Round B reaction conditions were as follows: 5 μL of Round A-labeled cDNA was added to 45 μL of KlenTaq master mix per sample (5 μL 10X KlenTaq PCR buffer, 1 μL 12.5 mM dNTP, 1 μL 100 pmol/μl Sol-PrimerB (5’- GTTTCCCACTGGAGGATA-3’), 1 μL KlenTaq LA (Sigma-Aldrich, St Louis, MO), 37 μL ddH_2_O). Reaction conditions for the PCR were as follows: 94°C for 2 minutes; 25 cycles of 94°C for 30 sec, 50°C for 45 sec, 72°C for 60 sec; 72°C for 5 minutes.

### Preparation of nanopore sequencing libraries

Amplified cDNA from Round B was purified using AMPure XP beads (Beckman Coulter, Brea, CA), and 1 μg DNA was used as input into Oxford Nanopore Genomic DNA MAP-003 Kits (Chik1, Ebola1) and MAP-004 Kits (HepC1, Ebola2) for generation of MinION Oxford Nanopore-compatible libraries[9, 11]. Briefly, the steps include (1) addition of control lambda phage DNA, (2) end-repair with the NEBNext End Repair Module, (3) 1X AMPure purification, (4) dA-tailing with the NEBNext dA-tailing Module, (5) ligation to protein-linked adapters HP/AMP (Oxford Nanopore Technologies, Oxford, UK) using the NEBNext QuickLigation Module x 10 min at room temperature, (6) purification of ligated libraries using magnetic His-Tag Dynabeads (Life Technologies), and (7) elution in 25 μL buffer (Oxford Nanopore Technologies). Lambda phage DNA was not added during preparation of the Ebola2 sample library.

### Nanopore sequencing

Nanopore libraries were run on an Oxford Nanopore MinION flow cell after loading 150 μL sequencing mix (6 μL library, 3 μL fuel mix, 141 μL BP mix) per the manufacturer’s instructions. The Chik1 and Ebola1 samples were run consecutively on the same flow cell, with an interim wash performed using Wash-Kit-001 (Oxford Nanopore).

### Illumina sequencing

For the Chik1 and Ebola1 specimens, amplified round B cDNA was purified using AMPure XP beads (Beckman Coulter) and 2 ng used as input into the Nextera XT Kit (Illumina). After 13 cycles of amplification, Illumina library concentration and average fragment size were determined using the Agilent Bioanalyzer. Sequencing was performed on an Illumina MiSeq using 150 nucleotide (nt) single-end runs and analyzed using the SURPI computational pipeline (UCSF, CA, USA) [15].

### MetaPORE bioinformatics pipeline

We developed a custom bioinformatics pipeline for real-time pathogen identification and visualization from nanopore sequencing data (MetaPORE), available at https://github.com/chiulab/MetaPORE. The MetaPORE pipeline consists of a set of Linux shell scripts, Python programs, and JavaScript / HTML code, and was tested and run on an Ubuntu 14.10 computational server with 64 cores and 512 GB memory. Alternatively, MetaPORE was tested and run on a laptop (4 cores, 32GB RAM, Ubuntu 14.10) by restricting the identification database to viral sequences, rather than all of NCBI nt.

Raw FAST5/HDF files from the MinION instrument are base-called using the Metrichor 2D Basecalling v1.14 pipeline (Metrichor). The MetaPORE pipeline continually scans the Metrichor download directory for batch analysis of downloaded sequence reads. For each batch of files (collected every time 200 reads are downloaded in the download directory or ≥2 min of elapsed time), the 2D read or either the template or complement read, depending on which is of higher quality, is converted into a FASTQ file using HDF5 Tools (https://www.hdfgroup.org/HDF5/doc/RM/Tools.html). The *cutadapt* program is then used to trim Sol-PrimerB adapter sequences from the ends of the reads[19]. Next, the nucleotide BLAST (BLASTn) aligner is used to computationally subtract host reads [15, 20] aligning to the human fraction of the National Center for Biotechnology Information (NCBI) nucleotide collection database (NT database, downloaded March 2015) at word size 11 and e-value cutoff of 10^-5^. On our 64 core machine, the remaining, non-human reads are then aligned by BLASTn to the entire NCBI nt database using the same parameters. On a laptop, the non-human reads are aligned to the viral fraction of the NCBI nt database. Reads that hit this viral database are then aligned by BLASTn to NCBI nt. For each read, the single best hit by e-value is retained. The NCBI GenBank gene identifier assigned to the best hit is then annotated by taxonomic lookup of the corresponding lineage, family, genus, and species [15].

For real-time visualization of results, a graphical user interface was developed for the MetaPORE pipeline. A “live” taxonomic count table is displayed as a donut chart using the CanvasJS (http://canvasjs.com) graphics suite, with the chart refreshing every 30 sec (Supplementary Data, Movie 1). For each viral family, genus, and species, the “top hit” is chosen to be the reference sequence with the greatest number of aligned reads, with priority given to reference sequences in the following order: (1) complete genomes; (2) complete sequence, or (3) partial sequences / individual genes. Coverage maps are generated in MetaPORE by mapping all aligned reads at a given taxonomic level (species, genus, or family) to the ‘top hit” using LASTZ v1.02 [21], with interactive visualization provided using a custom web program that accesses the HighCharts JavaScript library (http://www.highcharts.com). A corresponding interactive pairwise identity plot is generated by using SAMtools [22] to calculate the consensus FASTA sequence from the coverage map, followed by pairwise 100 bp sliding-window comparisons of the consensus to the reference sequence using the BioPython implementation of the Needleman-Wunsch algorithm [23, 24]. For purposes of comparison, the MetaPORE pipeline was also run on a subset of 100,000 reads from parallel Illumina MiSeq data corresponding to the Chik1, Ebola1, and Ebola2 samples.

### Phylogenetic Analysis

The overall CHIKV phylogeny (Fig. 2E, inset) consisted of all 188 near-complete or complete genome CHIKV sequences available in the NCBI nt database as of March 2015. A subphylogeny including the MiSeq-and nanopore-sequenced Puerto Rico strain PR-S6 presented here and previously[14], as well as additional Caribbean CHIKV strains and other representative members of the Asian-Pacific clade, was also analyzed (Fig. 2E). The EBOV phylogeny (Fig. 3E) consisted of the newly MiSeq-and nanopore-sequenced Ebola strain Lomela-LokoliaB11 from the 2014 DRC outbreak [25] and other representative EBOV strains, including strains from the 2014-2015 West African outbreak[8, 26]. Sequences were aligned using the MAFFT algorithm[27], and phylogenetic trees were constructed using the MrBayes algorithm [28] in the Geneious software package [29].

**Figure 2.**
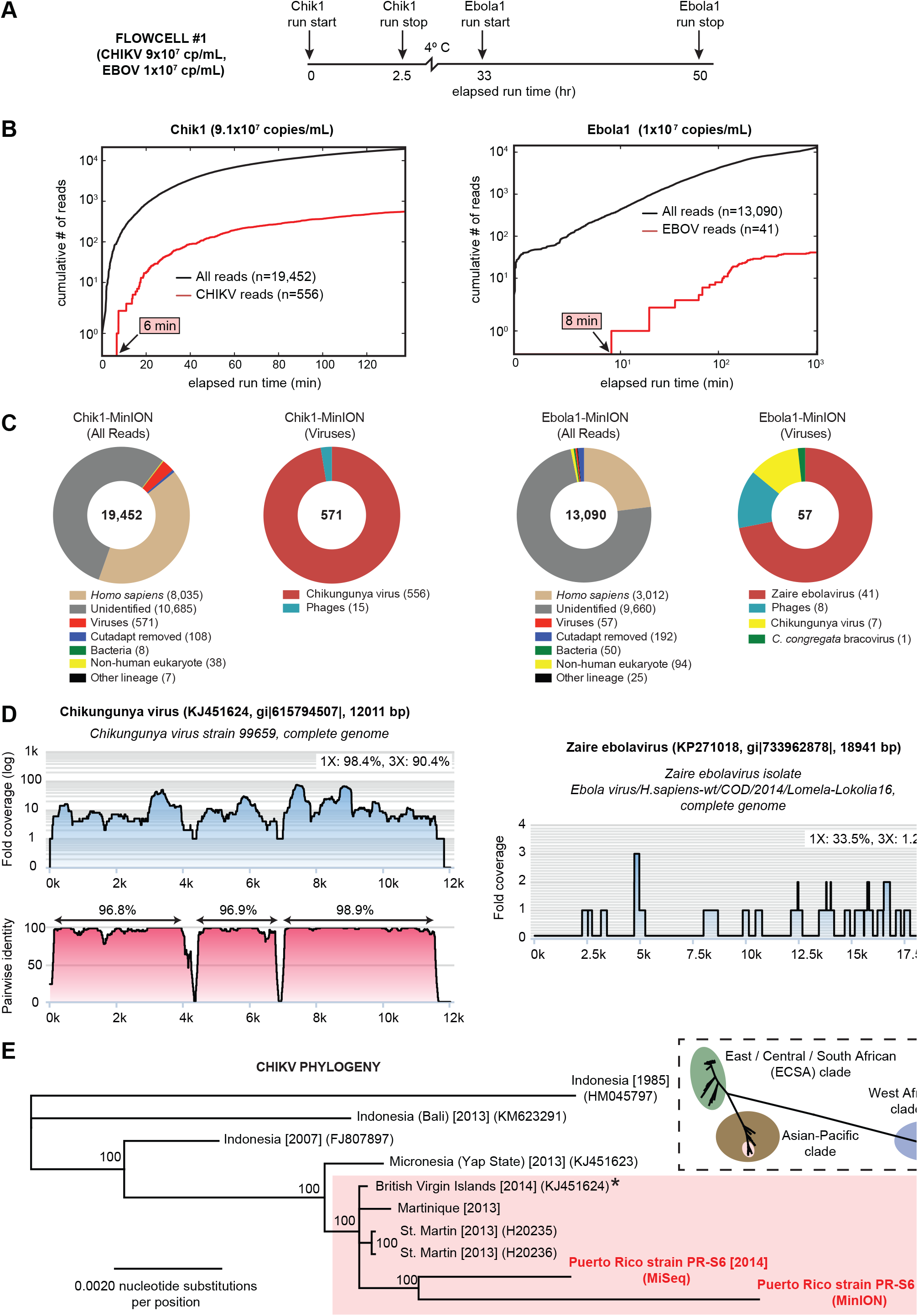
Metagenomic identification of CHIKV and EBOV from clinical blood samples by nanopore sequencing. **(A)** Timeline of sequencing runs on flowcell #1 with sample reloading, plotted as a function of elapsed time in hr since the start of flowcell sequencing. **(B)** Cumulative numbers of all sequenced reads (black line) and target viral reads (red line) from the Chik1 run (left) or Ebola1 run (right), plotted as a function of individual sequencing run time in min. **(C)** Taxonomic donut charts generated using the MetaPORE bioinformatics analysis pipeline from the Chik1 run (left) and Ebola1 run (right). The total number of reads analyzed is shown in the center of the donut. **(D)** Coverage and pairwise identity plots generated in MetaPORE by mapping reads aligning to CHIKV (left, Chik1 run) or EBOV (right, Ebola1 run) to the closest matching reference genome in the NCBI nt database (asterisk). **(E)** Whole-genome phylogeny of CHIKV. Representative CHIKV genome sequences from the Asian-Pacific clade, including the Puerto Rico PR-S6 strain recovered by nanopore and MiSeq sequencing, or all 188 near-complete or complete CHIKV genome sequences available in the NCBI nt database as of March 2015 (inset), are included. Branch lengths are drawn proportionally to the number of nucleotide substitutions per position, and support values are shown for each node. Abbreviations: CHIKV, chikungunya virus; EBOV, Ebola virus; Chik1, chikungunya virus, strain PR-S6 sample; Ebola1, EBOV, strain Lomela-Lokolia16 sample.

**Figure 3.**
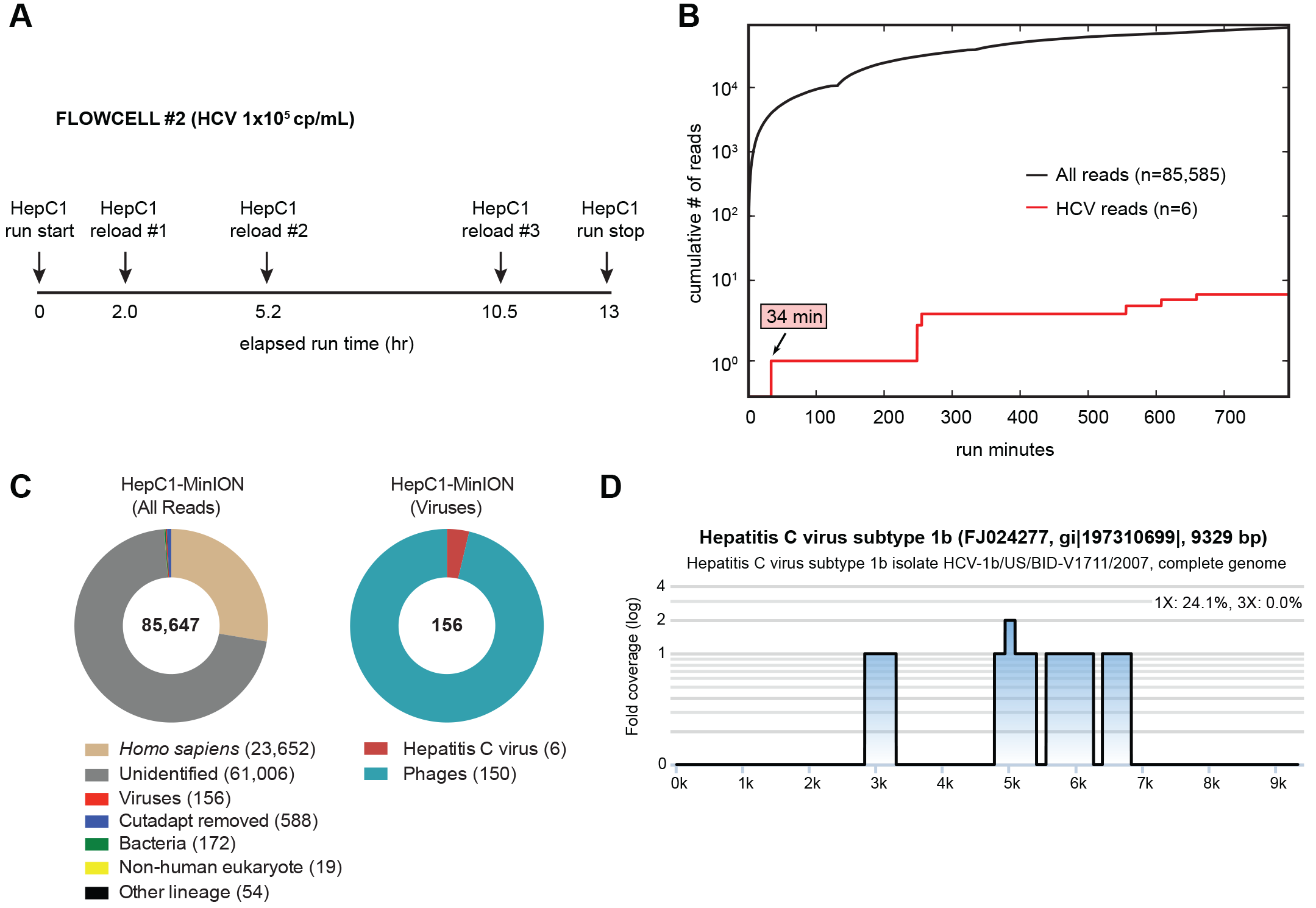
Metagenomic identification of HCV from a clinical serum sample by nanopore sequencing. **(A)** Timeline of sequencing runs on flowcell #2 with HepC1 sample reloading, plotted as a function of elapsed time in hr since the start of flowcell sequencing. **(B)** Cumulative number of all sequenced reads (black line) and HCV viral reads (red line), plotted as a function of individual sequencing run time in min. **(C)** Taxonomic donut charts generated using the MetaPORE bioinformatics analysis pipeline. The total number of reads analyzed is shown in the center of the donut. **(D)** Coverage and pairwise identity plots generated in MetaPORE by mapping reads aligning to HCV to the closest matching reference genome in the NCBI nt database. Abbreviations: HCV, hepatitis C virus; HepC1: hepatitis C virus, genotype 1b sample.

## RESULTS

### Example 1: High-titer CHIKV (Flowcell #1)

To test the ability of nanopore sequencing to identify metagenomic reads from a clinical sample, we first analyzed a plasma sample harboring high-titer CHIKV and previously sequenced on an Illumina MiSeq platform (Fig. 2B) [14]. The plasma sample corresponded to an asymptomatic blood donor who had screened positive for CHIKV infection during the 2014 outbreak in Puerto Rico (strain PR-S6), with a calculated viral titer of 9.1x10^7^ copies/mL.

A read aligning to CHIKV, the 96th read, was sequenced within 6 min (Fig. 2B) and detected by BLASTN alignment to the NCBI nt database within 8 min of data acquisition, demonstrating an overall sample-to-detection turnaround time of <6 hr (Fig. 1). After early termination of the sequencing run at the 2 hr 15 min time point, 556 of 19,452 total reads (2.85%) were found to align to CHIKV (Fig. 2B and C). The individual CHIKV nanopore reads had an average length of 455 nucleotides (nt) [range 126-1477] and average percent identity of 79% [range 51-92%] to the most closely matched reference strain, a CHIKV strain from the neighboring British Virgin Islands (KJ451624) (Table 1). When only high-quality 2D pass reads were included, 346 of 5,139 (6.7%) reads aligned to CHIKV, comparable to the proportion of CHIKV reads (248,677 of 3,235,096, 7.7%) identified by corresponding metagenomic sequencing on the Illumina MiSeq (Fig. S1).

Mapping of the 556 nanopore reads aligning to CHIKV to the assigned reference genome (KJ451624) showed recovery of 90% of the genome at 3X coverage and 98% at 1X coverage. Notably, despite high individual mapped error rates in the nanopore reads, ranging from 8 to 49%, 97-99% identity to the reference genome (KJ451624) was achieved across contiguous regions with at least 3X coverage (Fig. 2D). Furthermore, phylogenetic analysis revealed co-clustering of the CHIKV genomes independently assembled from MinION nanopore and Illumina MiSeq reads on the same branch within the Caribbean subclade (Fig. 2E). Overall, a large proportion of reads (55%) in the error-prone nanopore data remained unidentifiable, while other aligning reads aside from CHIKV corresponded to human, lambda phage control spike-in, uncultured bacterial, or other eukaryotic sequences (Fig. 2C).

### Example 2: High-titer Ebola virus (Flowcell #1)

We next attempted to replicate our metagenomic detection result on the nanopore sequencer with a different virus by testing a whole-blood sample from a patient with Ebola hemorrhagic fever during the August 2014 outbreak in the DRC (Ebola1, strain Lomela-Lokolia16) [25]. To conserve flowcells, the same nanopore flowcell used to run the Chik1 sample was washed and stored overnight at 4°C, followed by nanopore sequencing of the Ebola1 sample (viral titer of 1.0x10^7^ copies/mL by real-time qRT-PCR). Only 41 of 13,090 nanopore reads (0.31%) aligned to EBOV, as compared to 20,820 of 2,743,589 reads (0.76%) at 117X coverage for the Illumina MiSeq (Fig. 3B and D; Fig S1). The decrease in relative number (41 versus 556) and percentage (0.31% versus 2.85%) of target viral nanopore reads in the Ebola1 relative to Chik1 sample is consistent with the lower levels of viremia (1.0x10^7^ versus 9.1x10^7^ copies/mL) and high host background (whole blood versus plasma) (Fig. 2C). Nonetheless, the first read aligning to EBOV was detected in a similar timeframe as in the Chik1 sample, sequenced within 8 min and detected within 10 min of data acquisition. EBOV nanopore reads were 359 nt in length on average [range 220-672 nt], with a mean individual pairwise identity of 78% [range 56-88%]. Nevertheless, the majority of Ebola sequences (31 of 41, 76%) were found to align to the most closely matched strain of EBOV in the NCBI nt reference database, strain Lomela-Lokolia16 (Fig. 2D).

Despite washing the flowcell between the two successive runs, 7 CHIKV reads were recovered during the Ebola1 library sequencing, suggesting the potential for carryover contamination. CHIKV reads were not present in the corresponding Illumina MiSeq run (Fig. S1-B), confirming that the source of the contamination derived from the Chik1 library that was run on the same flow cell as the Ebola1 library.

### Example 3: Moderate-titer HCV (Flowcell #2)

Our previous experiments revealed both the total number of metagenomic reads and proportion of target viral reads at a given titer that could be obtained from a single MinION flowcell, and showed that the proportion of viral reads obtained by metagenomic nanopore and MiSeq sequencing was comparable. Thus, we projected that the minimum concentration of virus that could be reproducibly detected using our current metagenomic protocol would be 1x10^5^ copies/mL. An HCV-positive clinical sample (HepC1) was diluted in negative control serum matrix to a titer of 1x10^5^ copies/mL and processed for nanopore sequencing using an upgraded library preparation kit (MAP-004). After four consecutive runs on the same flowcell with repeat loading of the same metagenomic HepC1 library (Fig. 3A), a total of 85,647 reads were generated, of which only 6 (0.0070%) aligned to HCV (Fig. 3B). Although the entire series of flowcell runs lasted for >12 hr, the first HCV read was sequenced within 34 min (Fig. 3B), enabling BLASTn detection within 36 min of data acquisition. Given the low titer of HCV in the HepC1 sample and hence low corresponding fraction of HCV reads in the nanopore data, the vast majority (96%) of viral sequences identified corresponded to the background lambda phage spike-in. Importantly, although only 6 HCV reads were identified by nanopore sequencing, all 6 reads aligned to the correct genotype, genotype 1b (Fig. 3D).

### Example 4: High-titer Ebola virus with real-time MetaPORE analysis (Flowcell #3)

To enable real-time analysis of nanopore sequencing data, we combined our BLASTn analysis with monitoring and user-friendly web visualization into a real-time bioinformatics pipeline for pathogen detection named MetaPORE. We tested MetaPORE by sequencing a nanopore library (Ebola2) constructed using the upgraded MAP-004 kit and corresponding to a whole blood sample from a patient with suspected Ebola hemorrhagic fever during the 2014 DRC outbreak. Four consecutive runs of the Ebola2 library on the same flowcell over 34 hr, yielded a total of 335,044 reads, of which 593 (0.18%) aligned to EBOV (141 of 6009, or 2.3%, of 2D pass reads). Notably, the first EBOV read was sequenced 44 sec after data acquisition and correctly detected in ∼3 min by MetaPORE (Fig. 4B; Supplementary Information: Movie). A total of 3 EBOV reads, mapping across the genome and confirming unambiguous detection of the virus, were identified using MetaPORE within 9 min of data acquisition (Supplementary Information: Movie). In the corresponding Illumina MiSeq data, 1,778 reads (0.93%) out of a total of 192,158 reads aligned to EBOV by BLASTn, comparable to the proportion of EBOV reads in nanopore 2D pass data. Nanopore read coverage across the EBOV genome was relatively uniform with at least 1 read mapping to >88% of the genome and areas of zero coverage also seen with much higher-coverage Illumina MiSeq data (Fig. 4D). The detection of EBOV by real-time metagenomic nanopore sequencing was confirmed by qRT-PCR testing of the clinical blood sample, which was positive for EBOV at an estimated titer of 7.64 x10^7^ cp/ml. Phylogenetic analysis of the Ebola2 genome independently recovered by MinION nanopore and Illumina MiSeq sequencing revealed that nanopore sequencing alone was capable of pinpointing the correct EBOV outbreak strain and country of origin (Fig. 4E).

**Figure 4.**
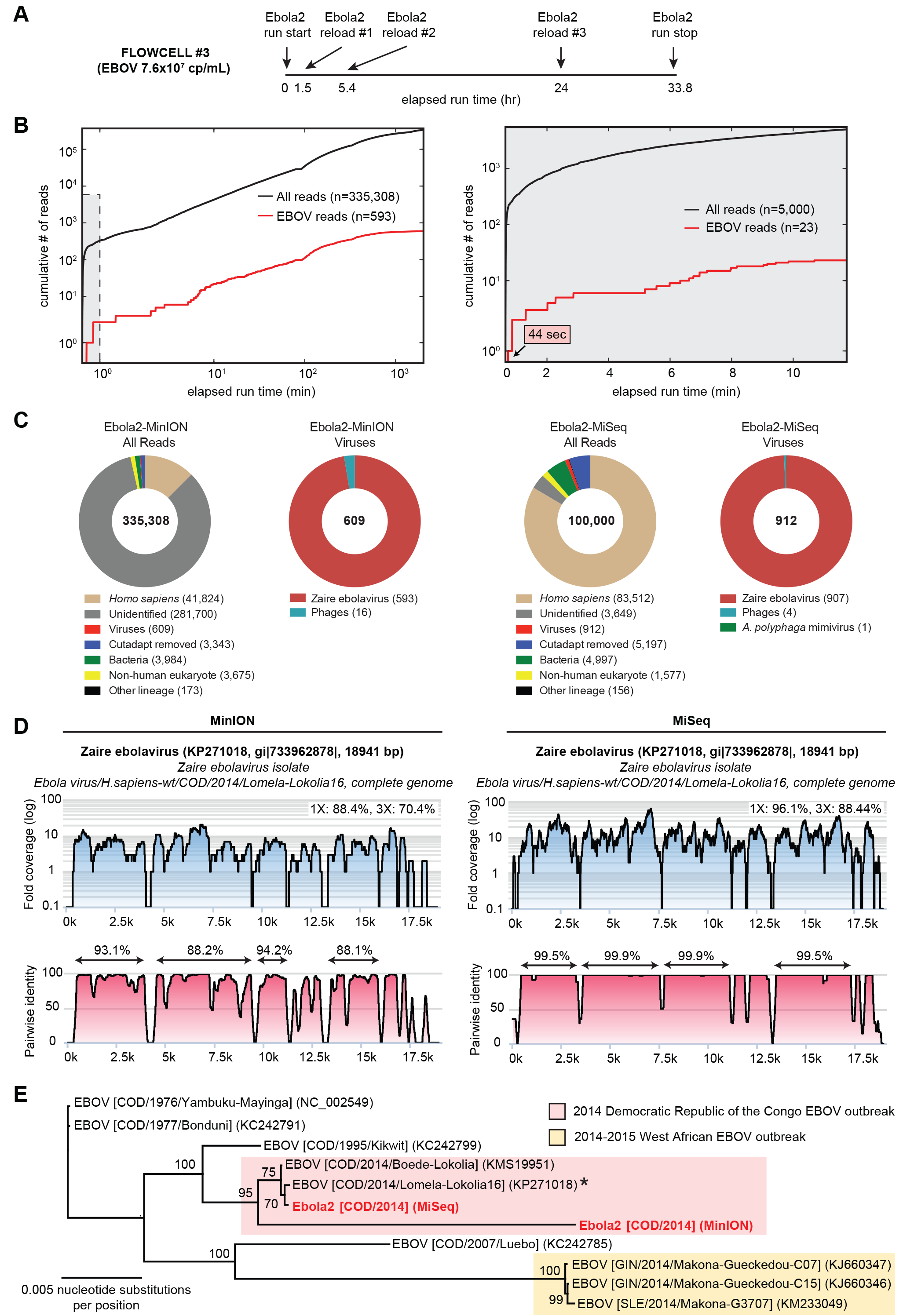
Metagenomic identification of EBOV from a clinical blood sample by nanopore sequencing and MetaPORE real-time bioinformatics analysis. Nanopore data generated from the Ebola2 library and sequenced on flowcell #3 was analyzed in real-time using the MetaPORE bioinformatics analysis pipeline, and compared to corresponding MiSeq data. **(A)** Timeline of nanopore sequencing runs on flowcell #3 with sample reloading, plotted as a function of elapsed time in hr since the start of flowcell sequencing. **(B)** Cumulative numbers of all sequenced reads (black line) and target viral reads (red line) from the nanopore run (left) or MiSeq run (right), plotted as a function of individual sequencing run time in min. **(C)** Taxonomic donut charts generated by real-time MetaPORE analysis of the nanopore reads (left) and post-run analysis of the MiSeq reads (right). The total number of reads analyzed is shown in the center of the donut. Note that only a subset of Illumina reads (n=100,000) were analyzed using the MetaPORE pipeline. **(D)** Coverage and pairwise identity plots generated by MetaPORE analysis of nanopore reads (left) and MiSeq reads (right), generated by mapping reads aligning to EBOV to the closest matching reference genome in the NCBI nt database (asterisk). Abbreviations: EBOV, Ebola virus; Ebola2, EBOV, strain Lomela-LokoliaB11 sample.

## DISCUSSION

Unbiased point-of-care testing for pathogens by rapid metagenomic sequencing has the potential to radically transform infectious disease diagnosis in clinical and public health settings. In this study, we sought to demonstrate the potential of the nanopore instrument for metagenomic pathogen identification in clinical samples by coupling an established assay protocol with a new, real-time sequence analysis pipeline. To date, high reported error rates (10-30%) and relatively low-throughput (<100,000 reads per flow cell) have hindered the utility of nanopore sequencing for analysis of metagenomic clinical samples [9, 11]. Prior work on infectious disease diagnostics using nanopore has focused on rapid PCR amplicon sequencing of viruses and bacteria[11], or real-time sequencing of pure bacterial isolates in culture, such *Salmonella* in a hospital outbreak [12]. To our knowledge, this is the first time that nanopore sequencing has been used for real-time metagenomic detection of pathogens in complex, high-background clinical samples in the setting of human infections. Here we also sequenced a microbial genome to high accuracy (>98% identity) directly from a metagenomic clinical sample and not from culture, using only multiple overlapping, albeit error-prone, nanopore reads and without resorting to the use of a secondary platform such as an Illumina MiSeq for sequence correction (Fig. 2D).

Real-time analysis is critical for time-critical sequencing applications such as outbreak investigation[7] and metagenomic diagnosis of life-threatening infections in hospitalized patients[3, 4, 6]. NGS analysis for clinical diagnostics is currently performed after sequencing is completed, analogous to how PCR products were analyzed by agarose gel electrophoresis in the 1990s. To date, most clinical PCR assays have been converted to a real-time format that reduces “hands-on” laboratory technician time and effort, decreases overall sample-to-answer turnaround times, and potentially yields quantitative information. Notably, our nanopore data suggest that very few reads are needed to provide an unambiguous diagnostic identification, despite high individual per read error rates of 10-30%. The ability of nanopore sequence analysis to accurate identify viruses to the species and even strain / genotype level is facilitated by the high specificity of sequence data, especially with the longer target viral reads achievable by nanopore (Table 1, 391 bp average length) versus second-generation sequencing.

Although the overall turnaround time from metagenomic sample-to-detection has been reduced to <6 hr, many challenges remain for routine implementation of this technology in clinical and public health settings. Improvements to make library preparation faster and more robust are critical, including automation and optimization of each step in the protocol. We also looked only at clinical samples at moderate to high titers of 10^5^-10^8^ copies / mL, and the sensitivity of metagenomic nanopore sequencing at lower titers remains unclear at current achievable sequencing depths. Standard wash protocols appear inadequate to prevent cross-contamination when reusing the same flow cell, as CHIKV reads were identified in the downstream Ebola1 sample sequence run. One solution may be to perform only one nanopore sequencing run per flow cell for clinical diagnostic purposes, akin to how disposable cartridges are used for clinical quantitative PCR testing on a Cepheid GenXpert instrument to prevent cross-contamination[30]. Another is to uniquely barcode individual samples at the cost of added time and effort. Finally, the current accuracy of a single nanopore read will most likely be insufficient to allow confident species identification of bacteria, fungi, or parasites, which have much larger genomes and shared conserved genomic regions than viruses. Single-nucleotide resolution will also be required for detection of antimicrobial resistance markers [31], which is difficult to achieve from relatively low-coverage metagenomic data [32]. These limitations can potentially be overcome by target enrichment methods such as capture probes to increase coverage, improvements in nanopore sequencing technology, or more accurate base-calling and alignment algorithms for nanopore data [33, 34].

## CONCLUSION

Our results indicate that unbiased metagenomic detection of viral pathogens from clinical samples with a sample-to-answer turnaround time of <6 hours and real-time bioinformatics analysis is feasible with nanopore sequencing. We demonstrate unbiased detection of diagnostic EBOV sequence in under four minutes after the Oxford Minion nanopore initiated sequencing. This technology will be particularly desirable for enabling point-of-care genomic analyses in the developing world, where critical resources, including reliable electric power, laboratory space, and computational server capacity, are often severely limited. MetaPORE, the real-time sequencing analysis platform developed here, is web-based and able to be run on a laptop. As sequencing yield, quality, and turnaround times continue to improve, we anticipate that third-generation technologies such as nanopore sequencing will challenge clinical diagnostic mainstays such as PCR and TMA testing, fulfilling the dream of an unbiased, point-of-care test for infectious diseases.

**Supplemental Figure 1. MetaPORE analysis of Illumina MiSeq data from samples containing CHIKV and EBOV. (A)** Taxonomic donut charts generated from MiSeq Chik1 run data (left) and Ebola1 run data (right) using the MetaPORE bioinformatics analysis pipeline. The total number of MiSeq reads analyzed is shown in the center of the donut. Note that only a subset of reads (n=100,000) were analyzed. **(D)** Coverage and pairwise identity plots generated in MetaPORE by mapping CHIKV reads from the Chik1 run (left) or EBOV reads from the Ebola1 run (right) to the closest matching reference genome in the NCBI nt database.

## Movie. MetaPORE Real-Time Bioinformatics Analysis and Visualization

(clip 0:19 - 10:13, 9.8 min) Detection of EBOV by metagenomic nanopore sequencing and real-time MetaPORE bioinformatics analysis. Raw FAST5 files are uploaded to the Metrichor cloud-based analytics platform for 2D basecalling (right panel). After downloading from Metrichor, basecalled FAST5 reads are collected in batches of 200 reads and automatically processed in real-time by MetaPORE. Reads corresponding to human host background are computationally subtracted by BLASTn alignment to the human portion of the NCBI nt database. Following subtraction, remaining reads are then aligned to NCBI nt using BLASTn for taxonomic identification. The closest matched reference genomes on the basis of hit frequency are shown in donut plots that are updated each minute in real-time (left panel). Note that the first EBOV read from the Ebola2 sequencing run is detected 3 min 6 sec (3:26) after the start of sequence acquisition (0:19). (clip 10:21 - 11:51, 1.5 min) Web-based, interactive coverage map and pairwise identity plots, generated in real-time by MetaPORE, enable zooming, highlighting of individual values, outputting of relevant statistical data, and exporting of the graphs in various formats. The plots shown in the movie correspond to the analyzed data after completion of nanopore sequencing.

## Data Availability

Nanopore and MiSeq sequencing data, along with sample metadata, have been submitted to NCBI under the following GenBank accession numbers: Sample metadata along with nanopore and MiSeq sequencing data have been submitted to the NCBI for each sample under the following SRA Study accessions: Ebola virus/H.sapiens-wt/COD/2014/Lomela-Lokolia16 (SRP057409), Ebola virus/H.sapiens-wt/COD/2014/Lomela-LokoliaB11 (SRS933322), Chik1 (SRP057410), and HepC1 (SRP057418).

## AUTHOR’S CONTRIBUTIONS

ALG, and CYC directed the study. ALG performed all nanopore experiments; SNN, GY, SS, and JB provided cDNA libraries and generated Illumina libraries. SF and ALG constructed the MetaPORE pipeline; SF,DS, CYC generated real time visualization; ALG, SF, SNN validated MetaPORE pipeline. VB, JML, RD, and SS provided chikungunya sample; PM, BSS, JJM provided ebola sample. ALG, SF, SNN, and CYC wrote the manuscript.

